# Disrupted intrinsic connectivity links to language and social deficits in toddlers with autism

**DOI:** 10.1101/2021.10.08.463640

**Authors:** Yaqiong Xiao, Teresa H. Wen, Lauren Kupis, Lisa T. Eyler, Disha Goel, Michael V. Lombardo, Karen Pierce, Eric Courchesne

## Abstract

Social and language abilities are closely intertwined during early development. Yet, it is still unknown how neural features underlying early social and language deficits are linked in toddlers with autism spectrum disorders (ASD). We examined functional connectivity of left and right temporal language regions and its correlations with language and social abilities in a cohort of 1– 4 years old toddlers (52 ASD/34 non-ASD). Further, ASD toddlers were stratified into those who strongly prefer social visual stimuli (ASD_Soc_) vs. those who do not (ASD_nonSoc_) based on performance on an eye-tracking paradigm. In non-ASD toddlers, connectivity between temporal regions and other language- and social-related cortical regions was significantly correlated with language, communication, and social scores. Conversely, ASD toddlers showed atypical correlations between temporal–visual cortex (cuneus) connectivity and communication ability. This temporal–visual connectivity was also correlated with social visual attention in ASD_nonSoc_ but not in ASD_Soc_ toddlers. These findings suggest language- and social-related functional connectivity was not correlated with language and social functions in ASD toddlers. Abnormal engagement of temporal–visual cortex connectivity may be an early-age signature of ASD and may help explain why interventions targeting social skills and language are so challenging, particularly in those with poor social engagement.

## Introduction

Early language and social communication deficits are often early warning signs of autism (Wetherby et al., 2004; Zwaigenbaum et al., 2005). Research with typically developing (TD) infants and toddlers has led to a theory that language and social development are closely intertwined in the early years of life (Kuhl, 2012, 2007). Specifically, social attention and social interactions are crucial for early language learning (Ferjan Ramírez et al., 2020; Kuhl, 2010; Ramírez-Esparza et al., 2017a, 2014). It is well recognized that typical language development is contingent on and constrained by early reciprocal social engagement (Klin et al., 2015; Kuhl, 2010), and that early language learning is associated with the emergence of the social brain in infants (Kuhl, 2010, 2007). Studies have reported that autistic children’s preference for affective speech (i.e., motherese) is linked to neural response to speech (Kuhl et al., 2005) and that neural response to known words with higher social skills predicts language outcomes at ages 4 and 6 in children with autism spectrum disorder (ASD) (Kuhl et al., 2013). Despite these findings reflecting important associations between language and social development in children with ASD, no studies have examined whether and how neural systems underlying language and social deficits are linked in early development of ASD.

It is well documented that left and right superior temporal regions are involved in language processing, especially in toddlers and young children (Holland et al., 2007; Olulade et al., 2020; Redcay et al., 2008). Dehaene-Lambertz and colleagues found robust activations in bilateral superior temporal regions during passive listening to speech in 2–3 month-old infants during natural sleep, and superior temporal cortex activation in infants was similar to that in adults (Dehaene-Lambertz et al., 2002). Research from our lab utilized a similar sleeping fMRI approach; fMRI data were collected from TD and ASD toddlers during passive listening to nursery stories during natural sleep. These several studies reproducibly report robust activation in superior temporal regions in TD toddlers but reduced superior temporal activation in ASD toddlers (Eyler et al., 2012; Redcay and Courchesne, 2008; Xiao et al., 2021). In addition, superior temporal activation to speech with varying levels of affect was related to a child’s social and communication abilities regardless of diagnosis (Xiao et al., 2021). In recent studies, we further reported that robust superior temporal activation to affective speech was observed not only in TD toddlers but also in ASD toddlers with good language outcomes, while reduced superior temporal activation occurred in ASD toddlers with poor language outcomes (Lombardo et al., 2015, 2018). Taken together, these findings demonstrate that reduced superior temporal activation in response to speech in different independent samples is a replicable, robust, and clinically relevant feature of deviant language emergence in the first years of life in ASD, especially in ASD toddlers with poor language outcomes.

Though still limited, a few ASD studies have revealed the neural basis of language processing and its development from a network perspective (Dinstein et al., 2011; Gao et al., 2019; Liu et al., 2020). For example, a functional connectivity study with ASD toddlers aged 1–4 years demonstrated poor inter-hemispheric connectivity in inferior frontal and superior temporal regions (Dinstein et al., 2011). Research also showed increased functional connectivity between language regions and between language and non-language regions (e.g., posterior cingulate cortex (PCC) and visual regions) in ASD vs. TD children and adolescents (ages 8–18 years) (Gao et al., 2019). These findings provide evidence for the neural bases of language processing beyond activation as they highlight ASD vs. TD differences in functional connectivity between language regions as well as between language and non-language regions. In addition, disrupted functional connectivity in ASD individuals are found to be behaviorally relevant (Dinstein et al., 2011; Gao et al., 2019). However, it is notable that in some at-risk infant studies (Liu et al., 2020), confirmed diagnoses were not reported, making it difficult to generalize at-risk findings to ASD. In fact, there are no studies that have investigated the relationship between language network connectivity and clinically behaviors (e.g., language and social abilities) in infants confirmed as ASD.

While language and social deficits are common symptoms in ASD (Lord et al., 2018, 2000; Wetherby et al., 2004), considerable heterogeneity exists across ASD individuals, and the early-age neural basis underlying this heterogeneity has been investigated in a few recent studies (Lombardo et al., 2019, 2015, 2018; Xiao et al., 2021). For example, two ASD subgroups were stratified based on language outcomes at 3–4 years, which showed strong associations with brain activation patterns in response to speech stimuli at 1–2 years (Lombardo et al., 2015, 2018). We have also discovered ASD subgroups based on visual attention preference for geometric over social images, as measured by an original eye tracking task known as the GeoPref test (Pierce et al., 2016b, 2011). The ASD subgroups identified by the GeoPref test not only differed in clinical scores (Lombardo et al., 2019; Pierce et al., 2016b, 2011) but also in functional connectivity between the default mode network (DMN) and the occipital-temporal cortex (Lombardo et al., 2019). Together, these findings suggest neural-behavior ASD subgroups can be identified on the basis of language ability or social visual preference. As language and social attention are highly intertwined in early development (Kuhl, 2012, 2010; Ramírez-Esparza et al., 2017a, 2014), one open question is whether ASD subgroups that display different social preferences also differ in neural correlates of language and social abilities.

Here, we collected resting-state fMRI data from a cohort of 86 ASD and non-ASD toddlers and examined functional connectivity of left and right superior temporal regions. As left and right superior temporal regions are engaged in both language and social cognitive processing (Deen et al., 2015; Kotz and Paulmann, 2011), we expected to see a widespread connectivity map including other language regions and regions related to social cognitive processing, such as the DMN regions (Mars et al., 2012; Schilbach et al., 2008). However, according to the interactive specialization theory that posits brain functional networks become specialized with development (Johnson, 2011, 2000), this extensive language network might be only present in toddlers (Liu et al., 2020) but not in adults. To test this hypothesis, we additionally collected resting-state fMRI data from awake TD adults. We also expected to see group differences in connectivity between ASD and non-ASD toddlers. Finally, we investigated the relevance of functional networks of left and right superior temporal regions with a toddler’s language, communication, and social skills. Since left superior temporal cortex is implicated in processing and comprehending the meaning of speech, we expected that its functional connectivity would be correlated with language scores. In contrast, functional connectivity of right superior temporal cortex would be linked to social and communication scores, since it engages in the processing of prosody in speech or music (Dehaene-Lambertz et al., 2010; Homae et al., 2006). We expected that these patterns are only in non-ASD toddlers. In contrast, there might be atypical brain–behavior correlations patterns in ASD toddlers, as a reflection of the linkage between language and social deficits and altered brain functional connectivity. We further examined the relations between network connectivity and social visual attention as measured by the GeoPref eye-tracking test, and expected distinct correlation patterns among ASD toddlers with stronger preference for and attention to nonsocial stimuli (ASD_nonSoc_) compared to toddlers with stronger preference for social stimuli (ASD_Soc_).

## Results

### Functional connectivity patterns in toddlers and adults

In both ASD and non-ASD toddlers, the functional connectivity analysis revealed a wide range of regions in the temporal and frontal cortices that showed significant correlations with left and right temporal ROIs (**Figure 1**). As expected, large swathes of lateral frontal and temporal cortex that highly overlap with language-relevant neural circuitry were prevalent. Additionally, along the medial surface of the brain, there is presence of important social-brain areas (i.e., DMN regions) such as anterior cingulate cortex (ACC), prefrontal cortex, lateral parietal lobe (LP), and PCC. Inferior temporal regions sensitive to face and object processing were also prevalent in connectivity maps. However, these DMN regions were not present in the functional connectivity maps in adults (**Figure 1**). No significant group differences (non-ASD vs. ASD) in iFC of left or right temporal ROI were observed after correcting for multiple comparison.

**Figure 1.**
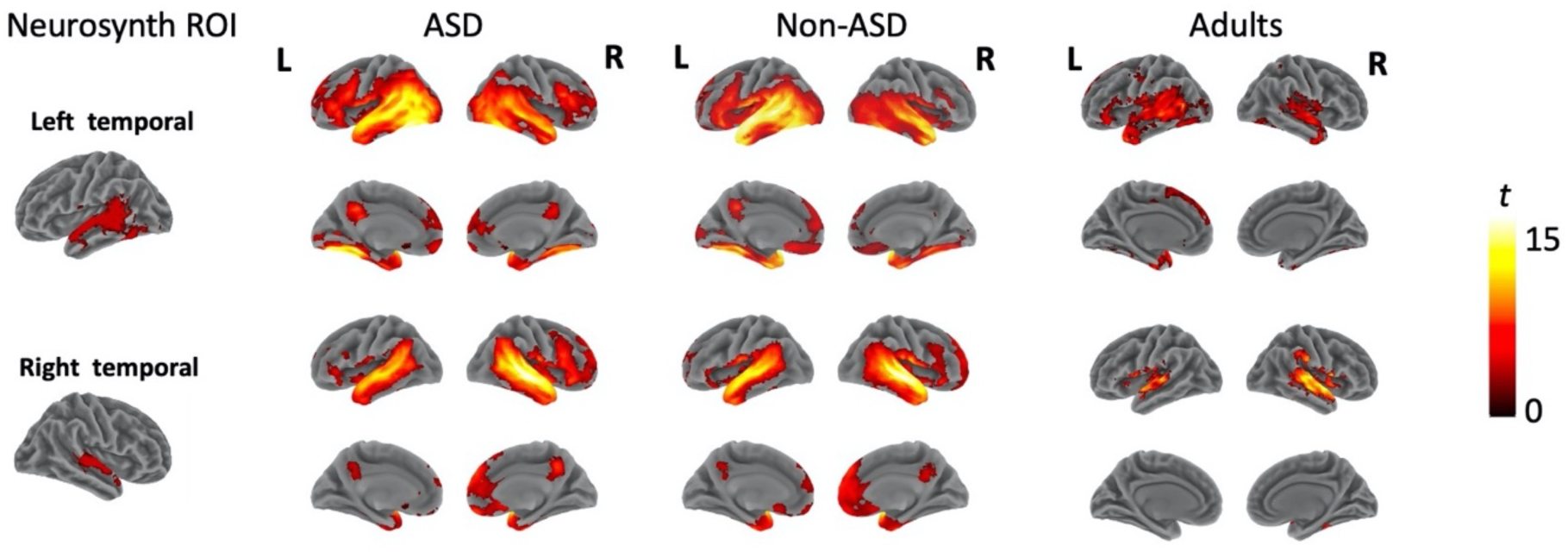
Whole-brain intrinsic functional connectivity maps for ASD, non-ASD, and adults. Results were corrected for multiple comparisons with voxel-wise *p* = 0.005 and cluster size > 155 voxels for toddlers and cluster size > 136 voxels for adults (cluster-wise *p* < 0.05, FWE corrected). ROI, region of interest; Abbreviations: ROI, region of interest; ASD, autism spectrum disorder; L, left; R, right.

### Connectivity–behavior correlation results

In non-ASD toddlers, we found that the strength of connectivity between temporal ROIs and other language and social regions was significantly correlated with language, communication, and social scores (**Figure 2 and Supplementary Table 1**). For the left temporal ROI, iFC with bilateral fronto-parietal operculum (FPO) was correlated with Mullen expressive language scores (**Figure 2A**). For the right temporal ROI, iFC with ACC and right LP was correlated with Mullen expressive language (**Figure 2C**), and iFC with ACC, right LP, and cerebellum was each correlated with Vineland communication (**Figure 2D**) and socialization scores (**Figure 2E**).

**Figure 2.**
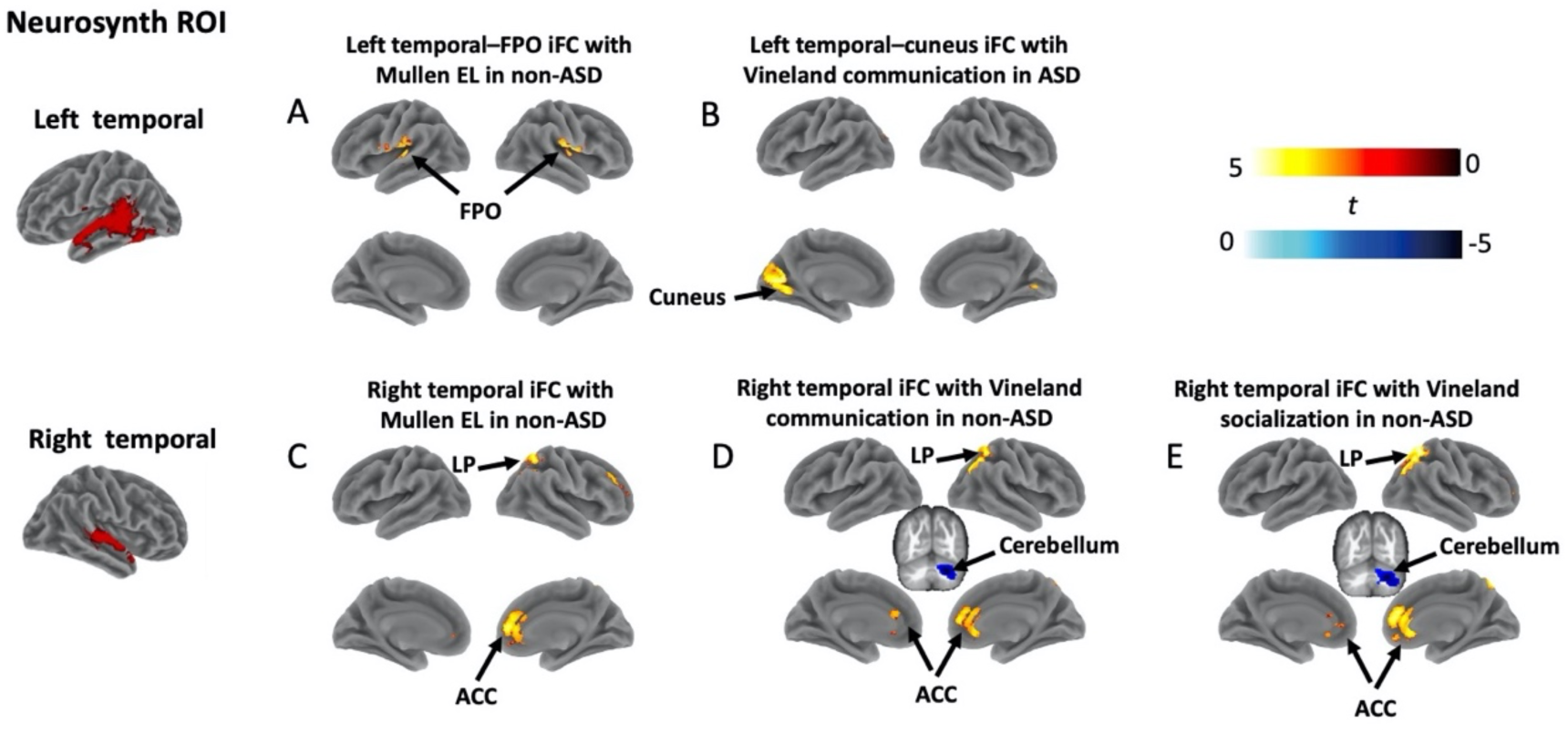
Clusters showing significant results in the connectivity–behavior correlation analysis. (A) iFC of left temporal ROI and bilateral fronto-periatal operculum was correlated with Mullen expressive language scores in non-ASD; (B) iFC of left temporal ROI and left cuneus linked to Vineland communications scores in ASD; (C) iFC of right temporal ROI and both ACC and LP linked to Mullen expressive language scores in non-ASD; (D) iFC of right temporal ROI and ACC, LP, and cerebellum linked to Vineland communication scores in non-ASD; (E) iFC of right temporal ROI and ACC, LP, and cerebellum linked to Vineland socialization scores in non-ASD. Clusters were corrected for multiple comparisons with voxel-wise *p* = 0.005 and cluster size > 155 voxels (cluster-wise *p* < 0.05, FWE corrected). Abbreviations: ROI, region of interest; ASD, autism spectrum disorder; iFC, intrinsic functional connectivity; EL, expressive language; FPO, fronto-parietal operculum; ACC, anterior cingulate cortex; LP, lateral parietal lobe.

These significant relationships in non-ASD toddlers were not found in ASD toddlers. Instead, we found that iFC between left temporal ROI and left cuneus was correlated with Vineland communication scores (**Figure 2B**), a relationship that was not present in non-ASD toddlers. Thus, ASD and non-ASD toddlers have distinctly different iFC patterns supporting language and social behaviors.

Further, we examined ASD vs. non-ASD group differences in correlation strength of iFC with clinical scores (see results in **Figure 3 and Supplementary Table 1**). For Mullen expressive language scores, there were significant group differences for left temporal ROI connectivity with bilateral FPO for expressive language scores (**Figure 3A**), and with left FPO for receptive language scores (**Figure 3B**). Also, there were significant group differences for right temporal ROI connectivity with right LP for Mullen expressive language scores (**Figure 3C**). For the Vineland, there were significant group differences for right temporal ROI connectivity with right LP for Vineland communication scores (**Figure 3D**), and with LP and cerebellum for Vineland socialization scores (**Figure 3E**).

**Figure 3.**
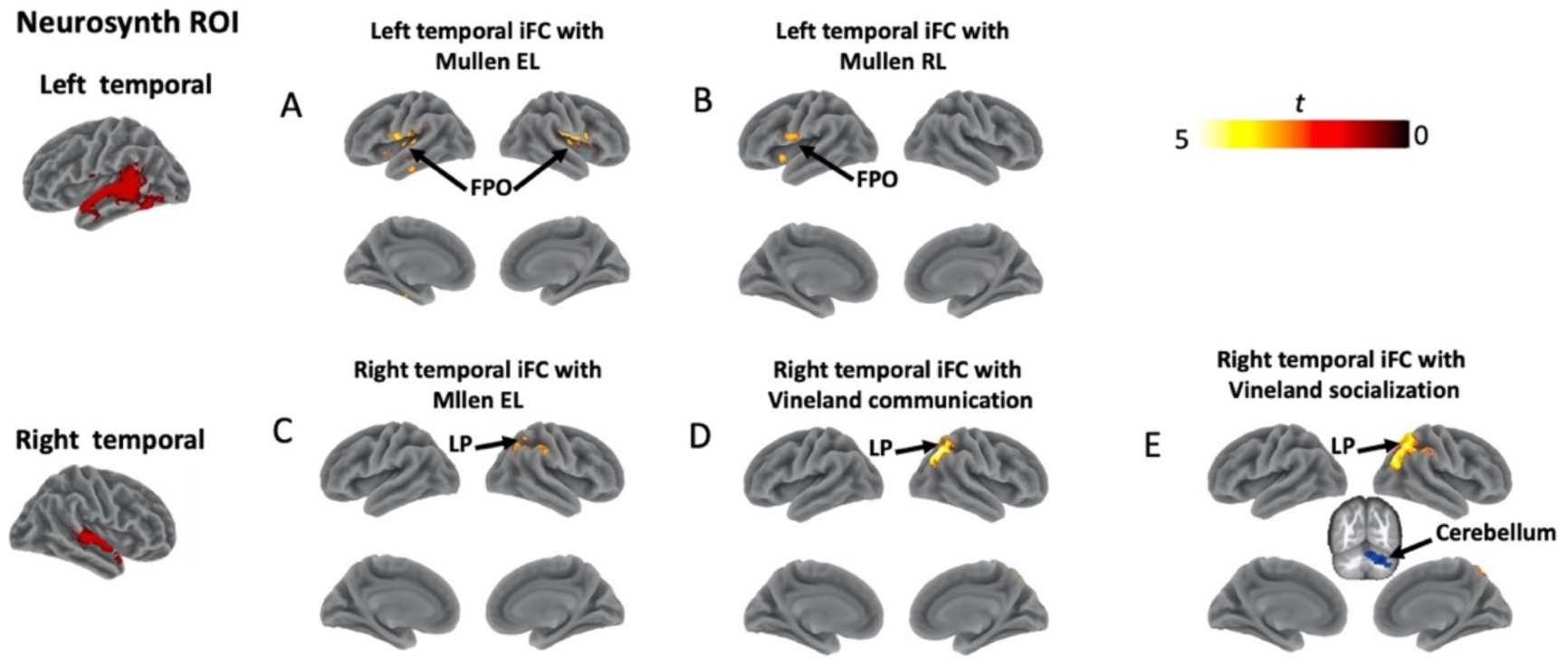
Clusters showing significant differences in brain-behavior correlations between non-ASD and ASD toddlers. Non-ASD and ASD groups differed significantly in correlations: (A) between iFC of left temporal ROI and bilateral FPO and Mullen expressive language scores; (B) between iFC of left temporal ROI and left FPO and Mullen receptive language scores; (C) between iFC of right temporal ROI and right LP and Mullen expressive language scores; (D) between iFC of right temporal ROI and right LP and Vineland communication scores; (E) between iFC of right temporal ROI and both right LP and cerebellum and Vineland socialization scores. Clusters were corrected for multiple comparisons with voxel-wise *p* = 0.005 and cluster size > 155 voxels (cluster-wise *p* < 0.05, FWE corrected). Abbreviations: ROI, region of interest; iFC, intrinsic functional connectivity; EL, expressive language; RL, receptive language; FPO, fronto-parietal operculum; LP, lateral parietal lobe.

### Subgroup-specific relationships between iFC and social and language abilities in ASD toddlers

Among 86 toddlers, 65 of them (33 ASD/32 non-ASD) had usable data from the eye-tracking test, and 33 ASD toddlers were designated as either nonsocial visual responders (ASD_nonSoc_: n = 16) or social visual responders (ASD_Soc_: n = 17) based on a threshold of 69% fixation to dynamic social stimuli (Pierce et al., 2016b) (**Figure 4A**). Specifically, toddlers were categorized as ASD nonsocial visual responders if they spent <69% of the duration of the video looking at social images, and toddlers were categorized as ASD social visual responders if they spent ≥69% of the duration of the video looking at dynamic social images. Using the same threshold, we identified 25 non-ASD toddlers as social visual responders and 7 as nonsocial visual responders, which were treated as a group (**Figure 4B**).

**Figure 4.**
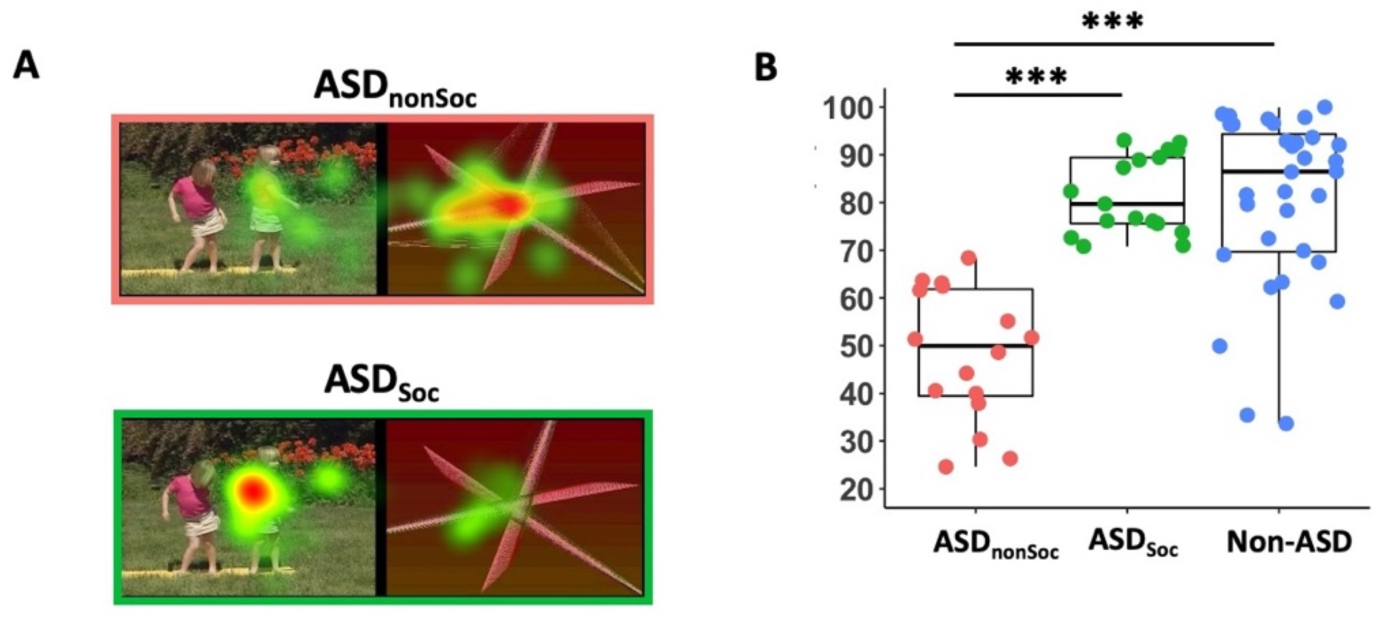
Identification of nonsocial and social visual ASD subgroups. (A) Examples of the stimuli used in the GeoPref eye-tracking test; example fixation from a nonsocial visual ASD (ASD_nonSoc_) individual (pink) and a social visual ASD (ASD_Soc_) individual (green). (B) Scatter-boxplot of GeoPref test performance for toddlers who also had resting-state fMRI data (ASD_nonSoc_: n = 16, pink; ASD_Soc_: n = 17, green; non-ASD: n = 32, blue). *** *p* < .001. Abbreviations: ASD, autism spectrum disorder.

We tested whether ASD_Soc_ and ASD_nonSoc_ subgroups differed in correlations between iFC of left and right temporal ROIs and language, communication, and social abilities. Functional connectivity values were extracted from cortical sites that had significant correlations in ASD or non-ASD or had significant ASD vs. non-ASD group differences for all toddlers (excluding repeated time points) (**Figures 2 and 3**).

Non-ASD toddlers have multiple significant positive correlations between left and right temporal ROIs and language, communication, and social scores ranging from 0.37 to 0.6 (**Supplementary Figure 1**). For the left temporal ROI in non-ASD, these included significant connectivity strength with bilateral FPO regions for Mullen expressive and receptive language scores (**Supplementary Figures 1B–C**). For the right temporal ROI in non-ASD, these included significant connectivity strength with the ACC and right LP for Mullen expressive language and Vineland communication scores (**Supplementary Figures 1E–H**). Moreover, right temporal ROI connectivity with ACC and LP was positively correlated with Vineland socialization scores (**Supplementary Figures 1J–K**). Interestingly, the only exception to positive correlations between temporal ROI iFC and language and social scores in non-ASD, were negative correlations involving the right temporal ROI connection with the cerebellum for Vineland socialization and communication scores.

Neither the ASD_Soc_ nor ASD_nonSoc_ subgroup had these positive correlations between temporal ROI iFC and language or social scores (**Supplementary Table 2**). Instead, ASD_Soc_ and ASD_nonSoc_ toddlers tended to display negative relationships, some being significantly negative. Remarkably, the only significantly positive connectivity–behavior relationship was one not present in non-ASD toddlers; namely, in both ASD_Soc_ and ASD_nonSoc_ subgroups, greater left temporal ROI connectivity with visual cortex (cuneus) was positively correlated with greater communication scores **(Supplementary Figure 1D)**.

Further, we observed significant subgroup differences (ASD_Soc_ vs. ASD_nonSoc_) in strength of correlation between right temporal ROI–right LP iFC and Mullen expressive language scores (*z* = 1.66, *p* = 0.049, one-tailed; **Supplementary Table 2**) and between right temporal ROI–cerebellum iFC and Vineland socialization scores (*z* = 1.95, *p* = 0.03, one-tailed; **Supplementary Table 2**). No other subgroup comparisons showed significant differences (**Supplementary Table 2**).

### Subgroup–specific associations between functional connectivity and ASD toddler’s social visual attention

There were 6 clusters (i.e., cuneus, left and right FPO, ACC, LP, and cerebellum) that were yielded by overlapping cortical sites that showed significant correlations in non-ASD or ASD or significant non-ASD vs. ASD group differences (**Figures 2 and 3**). We found subgroup differences (ASD_Soc_ vs. ASD_nonSoc_) in strength of correlation between left temporal ROI–cuneus and social visual attention differed (z = 1.94, *p* = 0.026, one-sided; **Figure 5A**), with a strong positive correlation in ASD_nonSoc_ (*r*(14) = 0.67, *p* = 0.004, 95% CI = [0.3, 0.87]) but no correlation in ASD_Soc_ (*r*(15) = 0.07, *p* = 0.79, 95% CI = [-0.39, 0.51]). We also observed strength of correlation between left temporal ROI–right FPO iFC and social visual attention differed between ASD subgroups (z = 1.78, *p* = 0.037, one-sided; **Figure 5B**), with a marginal negative correction in ASD_nonSoc_ (*r*(14) = −0.44, *p* = 0.086, 95% CI = [-0.79, −0.004]) and a pattern of positive correlation in ASD_Soc_ (*r*(15) = 0.21, *p* = 0.42 95% CI = [-0.28, 0.35]). No other subgroup differences (ASD_Soc_ vs. ASD_nonSoc_) in correlations between iFC and social visual attention were significant (**Figures 5C–F)**. For complete results of correlations in ASD subgroups as well as the comparisons between two ASD subgroups, see **Supplementary Table 3**.

**Figure 5.**
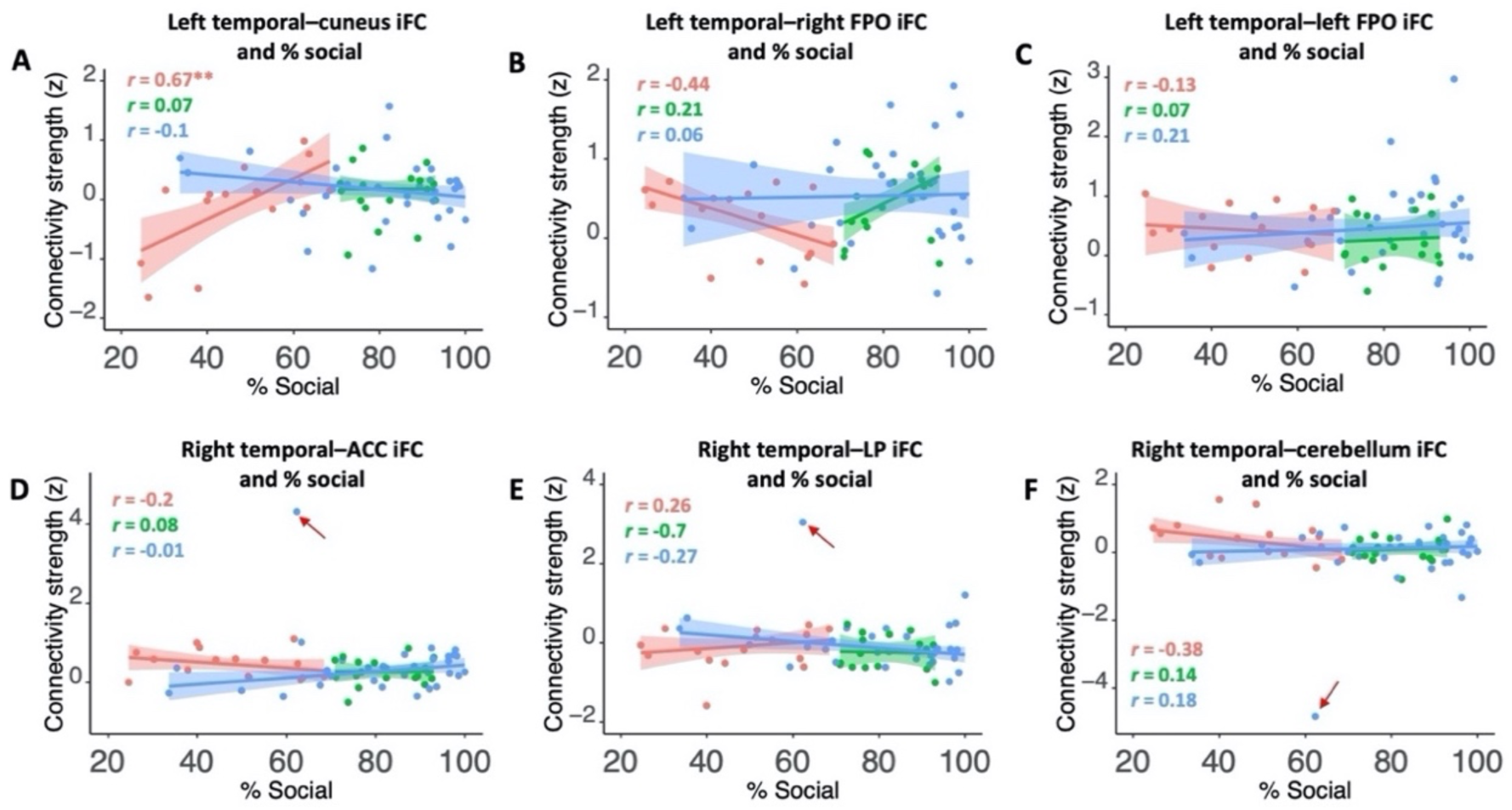
Scatterplots showing correlations between temporal iFC and social visual attention in ASD subgroups and non-ASD toddlers. Scatterplots display correlations for: (A) left temporal– cuneus iFC with % fixation on social images; (B) left temporal–left FPO iFC with % fixation on social images; (C) right temporal–right FPO iFC with % fixation on social images; (D) left temporal–ACC iFC with % fixation on social images (excluding the outlier in non-ASD: *r*(29) = 0.34; *p* = 0.06); (E) left temporal–LP iFC and % fixation on social images (excluding the outlier in non-ASD: *r*(29) = −0.2; *p* = 0.27); (F) left temporal–cerebellum iFC and % fixation on social images (excluding the outlier in non-ASD: *r*(29) = 0.02; *p* = 0.91). The relationships for ASD_nonSoc_ is shown in pink, for ASD_Soc_ shown in green, and for non-ASD shown in blue. The red arrows point to outliers. ** *p* < .01. Abbreviations: iFC, intrinsic functional connectivity; FPO, fronto-parietal operculum; ACC, anterior cingulate cortex; LP, lateral parietal lobe.

## Discussion

In non-ASD toddlers, we found that functional connectivity between temporal cortex ROIs and other language (FPO) and social regions (ACC, LP) was significantly correlated with clinical measures of language, communication, and social scores. These effects were absent in ASD toddlers. Thus, at early ages when language and social skills are rapidly developing, connectivity between language and social regions is not related to language and social functions in ASD toddlers; this brain–behavior deviance may help explain why interventions targeting social and language skills in ASD are so challenging. The most extreme deviation from non-ASD patterns was the presence of strong connectivity between language (i.e., left superior temporal) ROI and visual cortex (cuneus) linked to Vineland communication scores, something not present in non-ASD but a connection previously found in older ASD individuals (Gao et al., 2019; Kana et al., 2006; Koshino et al., 2005; Shen et al., 2012). This indicates that early-age neural network deviation may underpin how language is first acquired and processed in ASD toddlers across a range of ability levels. Importantly, the ASD_nonSoc_ subgroup had strong aberrant connectivity between left temporal ROI and visual cortex (cuneus) linked to low social visual attention. This pattern was not present in the ASD_Soc_ toddlers, who exhibited stronger social visual attention. Overall, in non-ASD toddlers, greater connectivity between temporal language cortex and language and social cortices (e.g., frontal, temporal, parietal, and anterior cingulate cortices) was correlated with better language, communication, and social scores, whereas ASD toddlers failed to even demonstrate trends towards positive correlations, instead showing near zero or slightly negative correlation trends.

Previous studies in TD toddlers consistently show language acquisition and development are closely linked to social experience (e.g., exposure to motherese, social responding, social interactions) in the first years of life (Goldstein et al., 2003; Goldstein and Schwade, 2008; Kuhl, 2007; Ramírez-Esparza et al., 2017a, 2017b, 2014). Here, our data demonstrate neural correlates (i.e., regions within language and social networks) of this in non-ASD toddlers. Specifically, in non-ASD, connectivity between the right temporal ROI and DMN regions (i.e., ACC and LP) was linked to language and social communication abilities. This finding provides neural evidence compatible with the view that language and social development are typically closely intertwined in the early years of life (Kuhl, 2012, 2007). It further suggests the hypothesis that social experience may promote language acquisition and learning and enhance functional connectivity between language and social brain regions as a result of the history of coactivation among these regions in non-ASD toddlers. In contrast, the lack of correlations between language-and social-related connectivity and language and social functions in ASD toddlers may reflect reduced social attention and social experience which leads to deviant language and social network connectivity inherent to ASD.

The atypical connectivity between left temporal cortex and cuneus in ASD toddlers in the present study has been consistently reported in previous studies of older ASD individuals (Gaffrey et al., 2007; Gao et al., 2019; Kana et al., 2006; Pang et al., 2016; Shen et al., 2012). In adult ASD vs. TD, Shen and colleagues (Shen et al., 2012) examined functional connectivity of the right extrastriate cortex, and found stronger connectivity between this seed region and bilateral frontal regions. Evidence also showed increased functional connectivity between language regions and visual cortex in children and adolescents with ASD vs. TD (Gao et al., 2019). These findings indicate the abnormal involvement of visual cortex in language processing in ASD children, adolescents, and adults. Here, in ASD toddlers ages 1–4 years, we found similar results, which provides evidence that this abnormal functional connectivity between language and visual regions may be present at a very early ages in this disorder.

Further, we observed different connectivity patterns underlying social visual attention between ASD_Soc_ vs. ASD_nonSoc_ subgroups, which has also been reported in our prior study (Lombardo et al., 2019). While the ASD_Soc_ subgroup, like non-ASD toddlers, showed almost no left temporal–cuneus connectivity correlation with social visual attention, the ASD_nonSoc_ subgroup had a strongly positive correlation. Notably, both ASD_Soc_ and ASD_nonSoc_ subgroups had positive correlations between temporal–cuneus connectivity and Vineland communication scores, a relationship that was absent in non-ASD toddlers. These findings are intriguing in two ways. First, for the ASD_nonSoc_ subgroup, correlations between temporal–cuneus connectivity and both communication and social visual attention suggest the possibility of aberrant developmental interactions between language and social domains at early ages (Kuhl, 2007; Ramírez-Esparza et al., 2017a, 2014). Second, ASD subgroups did not differ in correlations between temporal–cuneus connectivity and communication ability, indicating abnormal neural correlates of language deficits in both ASD subgroups. Thus, abnormal temporal–cuneus connectivity may be a more general signature of ASD. Future studies of social and language processing that examine the engagement of visual regions may shed light on this point.

One important finding is that our data show distinct brain–behavior correlation patterns for left and right temporal ROIs and for language and social abilities assessed by Mullen and Vineland subtests. Indeed, correlations of language and social network connectivity with language and social abilities are only present in the right temporal ROI, while the left temporal ROI shows only language ability correlations. These findings support that the left temporal region is implicated in language processing and comprehension, while the right superior temporal region is engaged in more emotionally and socially relevant features of communication. Further, the distinct correlation patterns for language and social subtests reflect the primary abilities that each subtest taps. For example, connectivity between left temporal ROI and bilateral FPO is only linked to Mullen expressive language but not Mullen receptive language scores. These findings suggest resting-state functional connectivity is a sensitive tool for detecting neural correlation patterns associated with different cognitive functions in the same domains (Uddin et al., 2010).

We observed different functional connectivity patterns in toddlers and adults. Early-age exuberant axonal connections have been well recognized (Huttenlocher, 1979; Innocenti and Price, 2005), which may explain widespread functional connectivity between temporal ROIs and other cortical regions involved in language and social processing in toddlers versus more constrained connectivity in adults. This finding also aligns with the interactive specification framework that argues the roles of brain regions gradually become specialized and restricted to a narrower set of functions with development (Johnson, 2011). Although resting-state data do not allow for direct inferences about specialization in response to tasks, it has been suggested that brain resting-state functional connectivity reflects recent experience and is also constrained by the underlying neuroanatomy (Uddin et al., 2010). Notably, in the present study, resting-state data were collected from sleeping toddlers and awake adults. However, research has shown consistent functional connectivity patterns in adults during light sleep and wakefulness (Larson-Prior et al., 2009). Thus, we expect the resting-state functional connectivity patterns are stable regardless of the status during the resting-state scan.

Contrast to our hypothesis, however, there were no significant differences in functional connectivity with temporal ROIs between ASD and non-ASD toddlers. Instead, we observed abnormal neural–behavior correlation patterns in ASD vs. non-ASD toddlers. It is possible that the abnormality of neural–behavior correlations in ASD toddlers is due to the underlying early-age axonal developmental defect. Indeed, we observed less deviant posterior structural connectivity in our previous study (Solso et al., 2016), which may additionally help explain why temporal–visual cortex connectivity stands out as a signal of strikingly aberrant functional language and social systems in the present study with young ASD toddlers as well as studies of older ASD individuals. Nevertheless, these findings suggest that, in order to gain a better understanding of what’s going awry in early years of ASD, it may be crucial to examine the connectivity–behavior correlations instead of just brain connectivity.

The present study has four potential limitations. First, we used a 2-year-old template for spatial normalization, which may not be optimal for all subjects, especially for those older children (>36 months). However, in this sample, 59 out of 86 (80%) subjects fell into the range of 1-3 years. Nevertheless, fMRI results could benefit from using a template that fits as many subjects as possible. Second, we did not analyze some contextual factors known to affect neurodevelopment including development in young children with ASD, such as family socioeconomic status (Olson et al., 2021a, 2021b) and language exposure at home. Future studies could consider these variables. Third, the sample size of ASD subgroups here is relatively small. Future work is necessary with larger samples in both subgroups. Nonetheless, we attempted to supplement the brain–behavior correlation estimates by reporting the bootstrap confidence intervals. Lastly, the distinct brain-behavior correlation patterns in non-ASD and ASD may be related to lower cognitive levels in ASD than non-ASD toddlers. It is possible that brain–behavior relationships observed in ASD toddlers are because they were not yet at the developmental stage where particular skills are required. Future research could include a mental-age-matched non-ASD group to control for the cognitive levels in non-ASD and ASD toddlers.

In conclusion, the present study revealed that functional connectivity within language regions and between language and social regions is related to language, communication, and social abilities in non-ASD but not ASD toddlers. Instead, ASD toddlers showed a correlation between temporo– visual cortex connectivity and communication ability, demonstrating abnormal and highly unusual involvement of this connectivity in communication deficits. Further, we observed that this temporo–cuneus connectivity differed between social and nonsocial ASD with regard to social visual attention. The absence of neurotypical brain-behavior correlations in language- and social-related regions coupled with the presence of the highly atypical engagement of temporo–cuneus connectivity may serve as a biomarker of early language and social deficits in ASD and may facilitate identifying ASD subgroups. Novel treatment approaches may be necessary to overcome and remodel these initial early-age, strikingly unusual neural networks for social and communication in ASD (Müller and Fishman, 2018).

## Materials and methods

### Participants

This study was approved by the University of California, San Diego Institutional Review Board. Informed consent was obtained from parents or guardians of toddlers, or from adult participants.

Applying an identical approach used in previous reports (Lombardo et al., 2019, 2015; Pierce et al., 2019, 2016a, 2011; Pramparo et al., 2015), we recruited toddlers through community referral and a population-based screening method in collaboration with pediatricians called *Get SET Early* (Pierce et al., 2021), which was formerly known as the 1-Year Well-Baby Check-Up Approach (Pierce et al., 2019, 2016a). Toddlers were assessed using the Autism Diagnostic Observation Schedule (ADOS-2; Module T, 1, or 2) (Lord et al., 2000), Mullen Scales of Early Learning(Mullen, 1995), and Vineland Adaptive Behavior Scales (Second Edition) (Sparrow et al., 2005). The Mullen assesses cognitive ability and development at ages 0 to 68 months; the Vineland assess adaptive communication, social, and behavioral functions. The diagnosis of ASD was based the DSM-5 diagnostic classification. Toddlers with initial diagnostic and clinical evaluations at < 36 months returned for follow-up evaluations. Clinical scores from a child’s most recent evaluation were used as the best estimate of abilities (**Table 1**). Assessments were administered by licensed, Ph.D.-level psychologists and occurred at the University of California, San Diego Autism Center of Excellence. Adult participants were recruited by word of mouth and had at least undergraduate education with no history of neurological, medical, or psychological disorders. Clinical, behavioral, and resting-state fMRI data were collected between 2018 and 2020 and have not been published before.

**Table 1.**
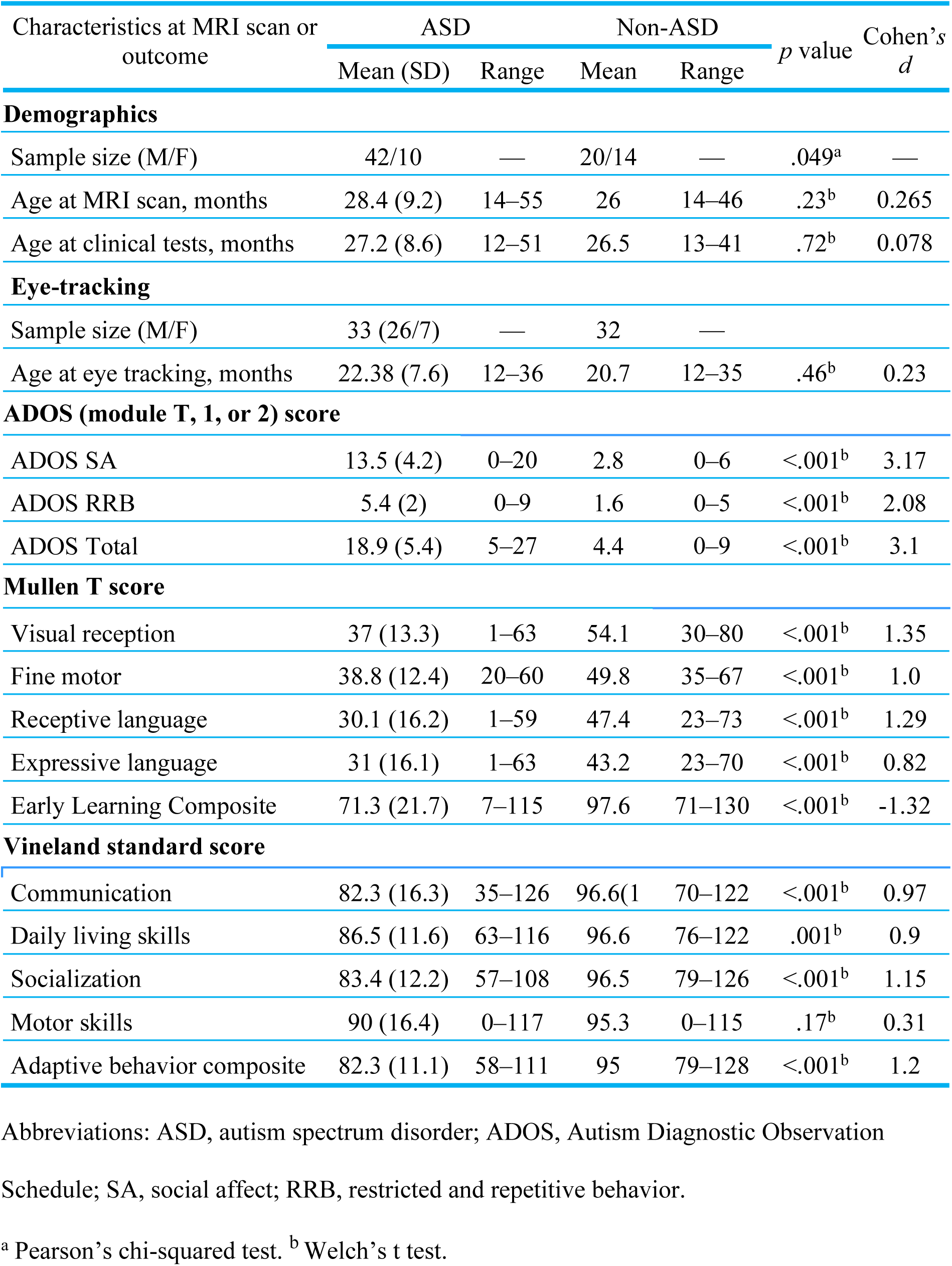
Demographic information and clinical test scores for ASD and non-ASD toddlers.

| Characteristics at MRI scan or outcome | ASD |  | Non-ASD |  | *p* value | Cohen's *d* |
| --- | --- | --- | --- | --- | --- | --- |
|  | Mean (SD) | Range | Mean | Range |  |  |
| Demographics |  |  |  |  |  |  |
| Sample size (M/F) | 42/10 | — | 20/14 | — | .049<sup>a</sup> | — |
| Age at MRI scan, months | 28.4 (9.2) | 14–55 | 26 | 14–46 | .23<sup>b</sup> | 0.265 |
| Age at clinical tests, months | 27.2 (8.6) | 12–51 | 26.5 | 13–41 | .72<sup>b</sup> | 0.078 |
| Eye-tracking |  |  |  |  |  |  |
| Sample size (M/F) | 33 (26/7) | — | 32 | — |  |  |
| Age at eye tracking, months | 22.38 (7.6) | 12–36 | 20.7 | 12–35 | .46<sup>b</sup> | 0.23 |
| ADOS (module T, 1, or 2) score |  |  |  |  |  |  |
| ADOS SA | 13.5 (4.2) | 0–20 | 2.8 | 0–6 | <.001<sup>b</sup> | 3.17 |
| ADOS RRB | 5.4 (2) | 0–9 | 1.6 | 0–5 | <.001<sup>b</sup> | 2.08 |
| ADOS Total | 18.9 (5.4) | 5–27 | 4.4 | 0–9 | <.001<sup>b</sup> | 3.1 |
| Mullen T score |  |  |  |  |  |  |
| Visual reception | 37 (13.3) | 1–63 | 54.1 | 30–80 | <.001<sup>b</sup> | 1.35 |
| Fine motor | 38.8 (12.4) | 20–60 | 49.8 | 35–67 | <.001<sup>b</sup> | 1.0 |
| Receptive language | 30.1 (16.2) | 1–59 | 47.4 | 23–73 | <.001<sup>b</sup> | 1.29 |
| Expressive language | 31 (16.1) | 1–63 | 43.2 | 23–70 | <.001<sup>b</sup> | 0.82 |
| Early Learning Composite | 71.3 (21.7) | 7–115 | 97.6 | 71–130 | <.001<sup>b</sup> | -1.32 |
| Vineland standard score |  |  |  |  |  |  |
| Communication | 82.3 (16.3) | 35–126 | 96.6(1 | 70–122 | <.001<sup>b</sup> | 0.97 |
| Daily living skills | 86.5 (11.6) | 63–116 | 96.6 | 76–122 | .001<sup>b</sup> | 0.9 |
| Socialization | 83.4 (12.2) | 57–108 | 96.5 | 79–126 | <.001<sup>b</sup> | 1.15 |
| Motor skills | 90 (16.4) | 0–117 | 95.3 | 0–115 | .17<sup>b</sup> | 0.31 |
| Adaptive behavior composite | 82.3 (11.1) | 58–111 | 95 | 79–128 | <.001<sup>b</sup> | 1.2 |

Abbreviations: ASD, autism spectrum disorder; ADOS, Autism Diagnostic Observation
Schedule; SA, social affect; RRB, restricted and repetitive behavior.
<sup>a</sup> Pearson's chi-squared test. <sup>b</sup> Welch's *t* test.

We collected resting-state fMRI data from N=86 naturally-sleeping toddlers ages 14 to 55 months (52 ASD/34 non-ASD) and from N=10 awake TD adults (6 F/4 M; 20–37 years old). Toddlers were considered non-ASD if their diagnosis at the outcome visit was not ASD and their Mullen Early Learning Composite score fell within 2 standard deviations of the mean score (i.e., > 70). This allowed us to examine brain functional connectivity along a continuum of language and cognitive abilities in non-ASD children. A subset of toddlers (5 ASD/5 non-ASD) had retest fMRI scans collected at intervals ranging from 2–22 months after the initial scan; 1 TD adult had a retest fMRI scan collected 20 days after the initial scan.

### GeoPref eye-tracking test

The 86 toddlers also participated in the eye-tracking GeoPref Test(Lombardo et al., 2019; Pierce et al., 2016a, 2011) wherein they watched two silent movies displaying dynamic geometric or dynamic social stimuli side by side. Dynamic geometric stimuli consisted of colourful moving geometric patterns, and dynamic social images consisted of children doing yoga exercises (**Figure 1A**). In order to control for spatial biases, spatial location of stimuli presentation (left/right) was randomly assigned across subjects. A total of 28 different geometric/social scenes of variable duration (0.8–3.7 seconds) were presented and lasted 62 seconds.

Eye tracking was conducted using Tobii software (Tobii Studio and Tobii Pro Lab), and fixation data were collected using a velocity threshold of 0.42 pixels/ms (Tobii Studio Tobii Fixation Filter) or 0.03 degrees/ms (Tobii Pro Lab Tobii IV-T Fixation Filter). Percent fixation to dynamic social images was used as an index of social visual attention and was computed by dividing fixation duration within an area of interest drawn around the dynamic social images by the total fixation duration across the entire video.

Of the total 86 toddlers with usable resting-state fMRI data, 65 of them (33 ASD/32 non-ASD) had moderate or good eye-tracking performance and total looking time > 50%. The other 21 toddlers either failed to complete the eye-tracking session, or the data quality was poor. Fifty-nine of these 65 toddlers completed the GeoPref test prior to the fMRI scan, and 6 toddlers completed the test after the fMRI scan.

### MRI data acquisition

Structural and functional MRI data were collected in a 3T GE scanner at the University of California, San Diego Center for Functional MRI. Resting-state functional images were acquired with a multi-echo EPI protocol (TE = 15ms, 28ms, 42ms, 56ms; TR = 2500ms; flip angle = 78°; matrix size = 64 × 64; slice thickness = 4 mm; field of view (FOV) = 256 mm; 34 slices, 288 volumes, a total of 12 minutes). Structural images were acquired using a T1-weighted MPRAGE sequence (FOV = 256 mm; TE = 3.172ms; TR= 8.142ms; Flip angle = 12°).

### Imaging data preprocessing

Multi-echo resting-state fMRI data were preprocessed using Multi-Echo Independent Components Analysis (ME-ICA) with a pipeline “meica.py” (ME-ICA 3.2) (Kundu et al. 2012, 2013) implemented in AFNI (Cox, 1996) and Python. Prior to preprocessing, the first 4 volumes were discarded to allow for magnetization to reach steady state. Preprocessing before data denoising included motion correction based on the first TE images (TE = 15 ms), slice timing correction for images of each TE, spatial normalization using an age-matched toddler template (i.e., 2-year-old template) (Shi et al., 2011) as the majority of the toddlers fell into this age range, and optimal combination of time series of all TEs. This optimal combination of multi-echo datasets has been empirically shown to considerably enhance temporal signal-to-noise ratio (tSNR) over single-echo EPI data (Kundu et al. 2013). Multi-echo principle component analysis (ME-PCA) was then applied as a dimensionality reduction technique before application of ME-ICA denoising. ME-ICA takes the ME-PCA reduced data and identifies independent components (ICs) that are then scored by pseudo-F statistics rho (ρ) and kappa (κ), which denote degree of non-BOLD and BOLD-related signal weightings based on TE-dependence analysis (Kundu et al., 2017, 2012). ICs with high rho (ρ) and low kappa (κ) are components of non-BOLD related signal and are removed as part of the denoising process, while ICs with low rho (ρ) and high kappa (κ) scores are components with high levels of BOLD-related signal and are retained. This ME-ICA denoising procedure has been shown to be effective at substantially increasing tSNR and successfully removes a large proportion of the head-motion related and other complex non-BOLD artifacts (Kundu et al., 2017, 2013, 2012; Lombardo et al., 2016; Power et al., 2018) and vastly improves test-retest reliability of functional connectivity measures (Lynch et al., 2020). The final multi-echo denoised data were used for subsequent seed-based connectivity analysis.

Head motion was quantified via framewise displacement (FD) (Power et al., 2012). The group average FD was minimal (mean FD < 0.11 mm) in toddlers (ASD: mean ± SD = 0.081 ± 0.036 mm, range 0.031–0.19 mm; non-ASD: mean ± SD = 0.11 ± 0.1 mm, range 0.038–0.58 mm) and adults (mean ± SD = 0.1 ± 0.056 mm, range 0.046–0.23 mm). There were no significant group differences between ASD and non-ASD toddlers (*t*(95) = 1.86, *p* = 0.07) or between toddlers and adults (ASD toddlers vs. adults: *t*(66) = −1.32, *p* = 0.21; non-ASD toddlers vs. adults: *t*(49) = 0.27, *p* = 0.79) in two-tailed two-sample *t*-tests, including all test and retest scans.

### Seed-based functional connectivity analysis

We selected two language-relevant regions of interest (ROIs), i.e., left and right temporal regions, from the meta-analytic activation map in Neurosynth (https://neurosynth.org/) with the term “language”. These ROIs were identical to those used in previous studies (Lombardo et al., 2015, 2018), and are displayed in **Figure 2 (first column)**. As these ROIs are from adult samples, they were co-registered to the age-matched toddler template using FSL’s flirt function (Jenkinson et al., 2002; Jenkinson and Smith, 2001).

For estimating seed-based functional connectivity, we used a multiple-echo independent component regression (ME-ICR) approach, which has been shown to be superior at estimating seed-based connectivity while adjusting for effective degrees of freedom based on the BOLD-dimensionality of each individual’s data (Kundu et al., 2017, 2013). Pearson’s correlation coefficients were computed on the “mefc” images, the ICs output from the ME-ICA pipeline. Computing seed-based connectivity based on this data has been shown to be a robust estimator of functional connectivity and allows for appropriate adjustment for effective degrees of freedom, denoted by the number of ICs, which vary from subject to subject (Kundu et al., 2013). Fisher *Z*-transformation were computed on the connectivity maps, and then a 6 mm FWHM smoothing kernel was applied to enhance SNR before group-level comparisons.

### Whole-brain group analysis

Considering that scans were collected at multiple time points, including test and retest scans, we ran whole-brain group analyses with mixed effects models using 3dMVM program (Chen et al., 2014) in AFNI (Cox, 1996) to take into account the repeated measures:

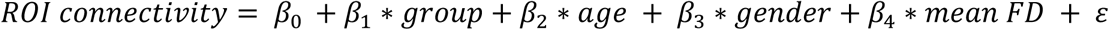

In separate models, iFC of left and right temporal ROIs served as a dependent variable, and subjects were treated as a random effect, which allows for fixed effects (i.e., age, sex, and mean FD) to vary for each subject.

Using a similar approach, we conducted the whole-brain group analysis with adult data. However, only within-group tests were performed as only TD adults were included.

Resulting correlation coefficient maps were corrected for multiple comparisons with the family-wise error (FWE) approach using 3dClustSim program in AFNI (voxel-wise *p* = 0.005 and cluster size > 136 voxels for adults and cluster size > 155 voxels for toddlers). This spatial cluster correction took into account spatial autocorrelation by using the ‘–acf’ option in 3dClustSim.

### Connectivity–behavior correlation analysis

For the main analysis, we investigated the relationships between iFC of language ROIs and a toddler’s language, communication, and social scores as assessed by the Mullen (Mullen expressive and receptive subscales) and Vineland (Vineland communication and socialization subscales). Mullen expressive and receptive language tap different aspects of language abilities, with expressive language tapping speaking and receptive language tapping auditory comprehension (Mullen, 1995). As expressive language is related to social skills (Pickard and Ingersoll, 2014; Schietecatte et al., 2012), Vineland communication which taps language abilities including both expressive and receptive language and written language is supposed to be an interaction of language and social constructs (Sparrow et al., 2005). Vineland socialization indexes social functioning within age-normed contexts.

For Mullen T scores, 21.2% of ASD toddlers (11 out of 52) in our study performed at levels that were below 20, and in order to enhance accuracy for brain-behavior correlations, we elected to generate a score for each toddler that was an approximate reflection of their ability, rather than artificially assigning all such toddlers a score of 20. The lowest minimum subscale T score based on the Mullen scoring manual is 20. Some ASD toddlers (i.e., 11 out of 52) in our study performed at levels that were below 20. For these cases, in order to enhance accuracy for brain-behavior correlations, we elected to generate a score for each toddler that was an approximate reflection of their ability, rather than artificially assigning all such toddlers a score of 20. Specifically, we estimated Mullen T-scores based on raw scores and a child’s chronological age using the following process: The estimated T score is calculated by examining the variation in raw scores for the lowest T scores available for the child’s age and applying that variation to estimate a lower T score. For example, if the lowest raw score available for the child’s age is 14, but the child actually has a raw score of 12, two steps would be counted down and an estimated score would be calculated based on the amount of difference between T scores for each raw score above the cut off. So, if the raw score of 14 corresponded to a T of 20, and there was a 2 point difference between each T score above 20, then the estimated T score would be 16 (2 steps times 2 point difference = 4 and thus the estimated T score is 20 - 4 = 16).

Similar to the whole-brain group analysis, we ran mixed effects models using 3dMVM program (Chen et al., 2014) in AFNI (Cox, 1996) with variables of interest (i.e., language, communication, and social scores) and those of no interest (i.e., age, gender, and mean FD) in the models. These behavioral variables included Mullen expressive and receptive language and Vineland social and communication scores which index toddlers’ language and social abilities (Sparrow et al., 2005; Mullen 1995). All resulting maps were transformed to *t*-maps. The same threshold as used for whole-brain group results (i.e., voxel-wise *p* = 0.005, cluster size > 155 voxels) was applied to the brain-behavior correlation coefficient maps.

Subsequently, we displayed the iFC–behavior correlation patterns (excluding repeated time points) and examined differences between ASD_Soc_ vs. ASD_nonSoc_ subgroups in correlation strength. First, we extracted *Z*-transformed correlation coefficients from clusters that had significant correlations with behaviors in non-ASD, ASD, or comparisons between non-ASD and ASD. Next, we presented scatterplots with trend lines for non-ASD group and ASD_Soc_, ASD_nonSoc_ subgroups. Finally, we tested group differences between ASD_Soc_ vs. ASD_nonSoc_ subgroups in the strength of correlations using the paired.r function in R (‘psych’ library) which computed *z*-statistics and *p*-values. Considering the relatively small sample sizes in two ASD subgroups, we performed bootstrapping to compute 95% confidence intervals (CI) around sample correlation estimates, using 100,000 bootstrap resamples, as this analysis allowed for reporting the distribution of sample correlation estimates that could have been observed.

Following similar procedures, we tested whether the relationships between functional connectivity and a toddler’s social visual attention—indexed by percent fixation to social images in the GeoPref test—differed between ASD_Soc_ and ASD_nonSoc_ subgroups. Specifically, functional connectivity values were extracted from overlapping cortical sites that showed significant correlations in non-ASD or ASD or significant ASD vs. non-ASD group differences. Then, we presented scatterplots with trend lines for non-ASD group and ASD_Soc_, ASD_nonSoc_ subgroups. Here, we included all non-ASD toddlers (n=32) in the scatterplots because the focus of the analysis was to compare the correlation strength between ASD_Soc_ vs. ASD_nonSoc_ subgroups (using paired.r function in R). Finally, we ran the same bootstrapping analysis as aforementioned to report the distribution of sample correlation estimates that could have been observed.

## Data availability

Tidy data are available at https://github.com/Yaqiongxiao/asdlanguage_rsfMRI.

## Acknowledgements

We are grateful to the parents and children in San Diego who participated in our research, without whom this would not be possible. We are also fortunate to work with wonderful pediatricians and family practice physicians spanning a range of medical groups including UCSD, Sharp Rees-Stealy, Scripps, Rady-Children’s Primary Care Medical Group, Chula Vista Pediatrics, Graybill Medical Group, Grossmont Pediatrics, Linda Vista Health Care Center, Mills Pediatrics, North County Health Services, San Diego Family Care, and Sea Breeze Pediatrics. We are grateful for their support.

## Supplementary tables and figures

**Supplementary Table 1.** Clusters showing significant connectivity–behavior relationships.

| Group/Contrast | Behavior (scores) | Region | MNI coordinates |  |  | Cluster size<br>(voxels) |
| --- | --- | --- | --- | --- | --- | --- |
|  |  |  | x | y | z |  |
| Left temporal ROI |  |  |  |  |  |  |
| ASD | Vineland | Cuneus | -15 | -67 | 3 | 312 |
| Non-ASD | Mullen EL | Left FPO | -45 | -20 | 3 | 199 |
|  |  | Right FPO | 45 | -14 | 9 | 184 |
| Non-ASD vs.<br>ASD | Mullen EL | left FPO | -42 | -20 | 6 | 378 |
|  |  | right FPO | 45 | -10 | 9 | 249 |
| Non-ASD vs. | Mullen RL | left FPO | -49 | 13 | -7 | 168 |
| Right temporal ROI |  |  |  |  |  |  |
| Non-ASD | Vineland<br>communication | Right LP | 32 | -60 | 40 | 310 |
|  |  | ACC | 8 | 40 | 16 | 241 |
|  |  | Cerebellum | 22 | -50 | -37 | 202 |
| Non-ASD vs. | Vineland | Right LP | 32 | -60 | 36 | 384 |
| Non-ASD | Vineland<br>socialization | Right LP | 29 | -71 | 26 | 485 |
|  |  | ACC | 8 | 40 | 19 | 343 |
|  |  | Cerebellum | 22 | -50 | -38 | 342 |
| Non-ASD vs.<br>ASD | Vineland<br>socialization | Right LP | 29 | -67 | 23 | 648 |
|  |  | Cerebellum | 22 | -50 | -38 | 214 |
| Non-ASD | Mullen EL | ACC | 8 | 40 | 16 | 332 |
|  |  | Right LP | 25 | -60 | 46 | 221 |
| Non-ASD vs. | Mullen EL | Right LP | 25 | -71 | 30 | 158 |

Clusters were corrected for multiple comparisons with voxel-wise $p = 0.005$ and cluster size $> 155$ voxels (cluster-wise $p < 0.05$ , FWE corrected). Abbreviations: EL, expressive language; RL, receptive language; ASD, autism spectrum disorder; FPO, fronto-parietal operculum; LP, lateral parietal cortex; ACC, anterior cingulate cortex.

**Supplementary Table 2.** Subgroup-specific relationships between iFC and social and language abilities as well as the comparisons between ASD subgroups.

| Behavior | Region | ASD<sub>nonSoc</sub> |  |  | ASD<sub>Soc</sub> |  |  | ASD<sub>Soc</sub> vs. ASD<sub>nonSoc</sub> |  |
| --- | --- | --- | --- | --- | --- | --- | --- | --- | --- |
|  |  | *r* value | *p* value | 95% CI | *r* value | *p* value | 95% CI | *z* value | *p* value |
| Mullen expressive language | Left FPO | -0.44 | 0.09 | [-0.73, -0.12] | -0.11 | 0.68 | [-0.47, 0.33] | 0.95 | 0.17 |
|  | Right FPO | -0.11 | 0.69 | [-0.57, 0.28] | -0.41 | 0.10 | [-0.78, 0.03] | 0.86 | 0.2 |
|  | ACC | 0.004 | 0.99 | [-0.44, 0.50] | -0.14 | 0.58 | [-0.62, 0.39] | 0.36 | 0.36 |
|  | Right LP | 0.16 | 0.57 | [-0.46, 0.73] | -0.45 | 0.07 | [-0.74, -0.001] | 1.66 | 0.049 |
| Mullen receptive language | Left FPO | -0.49 | 0.05 | [-0.75, 0.09] | -0.5 | 0.04 | [-0.76, -0.14] | 0.02 | 0.49 |
| Vineland communication | Cuneus | 0.58 | 0.02 | [0.13, 0.85] | 0.64 | 0.006 | [-0.05, 0.9] | 0.24 | 0.41 |
|  | Cerebellum | 0.07 | 0.80 | [-0.48, 0.67] | -0.21 | 0.42 | [-0.58, 0.18] | 0.73 | 0.23 |
|  | ACC | -0.23 | 0.39 | [-0.59, 0.32] | -0.36 | 0.16 | [-0.75, 0.46] | 0.37 | 0.36 |
|  | Right LP | -0.32 | 0.23 | [-0.66, 0.19] | -0.44 | 0.07 | [-0.79, 0.45] | 0.38 | 0.35 |
| Vineland socialization | ACC | -0.42 | 0.11 | [-0.71, 0.04] | -0.30 | 0.24 | [-0.75, 0.33] | 0.34 | 0.37 |
|  | Cerebellum | 0.40 | 0.13 | [-0.25, 0.74] | -0.32 | 0.21 | [-0.65, 0.10] | 1.95 | 0.03 |
|  | Right LP | -0.45 | 0.08 | [-0.78, -0.01] | -0.41 | 0.10 | [-0.81, 0.25] | 0.12 | 0.45 |

Abbreviations: ASD, autism spectrum disorder; ASDnonSoc, non-social visual responders with ASD; ASDSoc, social visual responders with ASD; CI, confidence intervals; ACC, anterior cingulate cortex; LP, lateral parietal lobe; FPO, fronto-parietal operculum.

**Supplementary Table 3.**
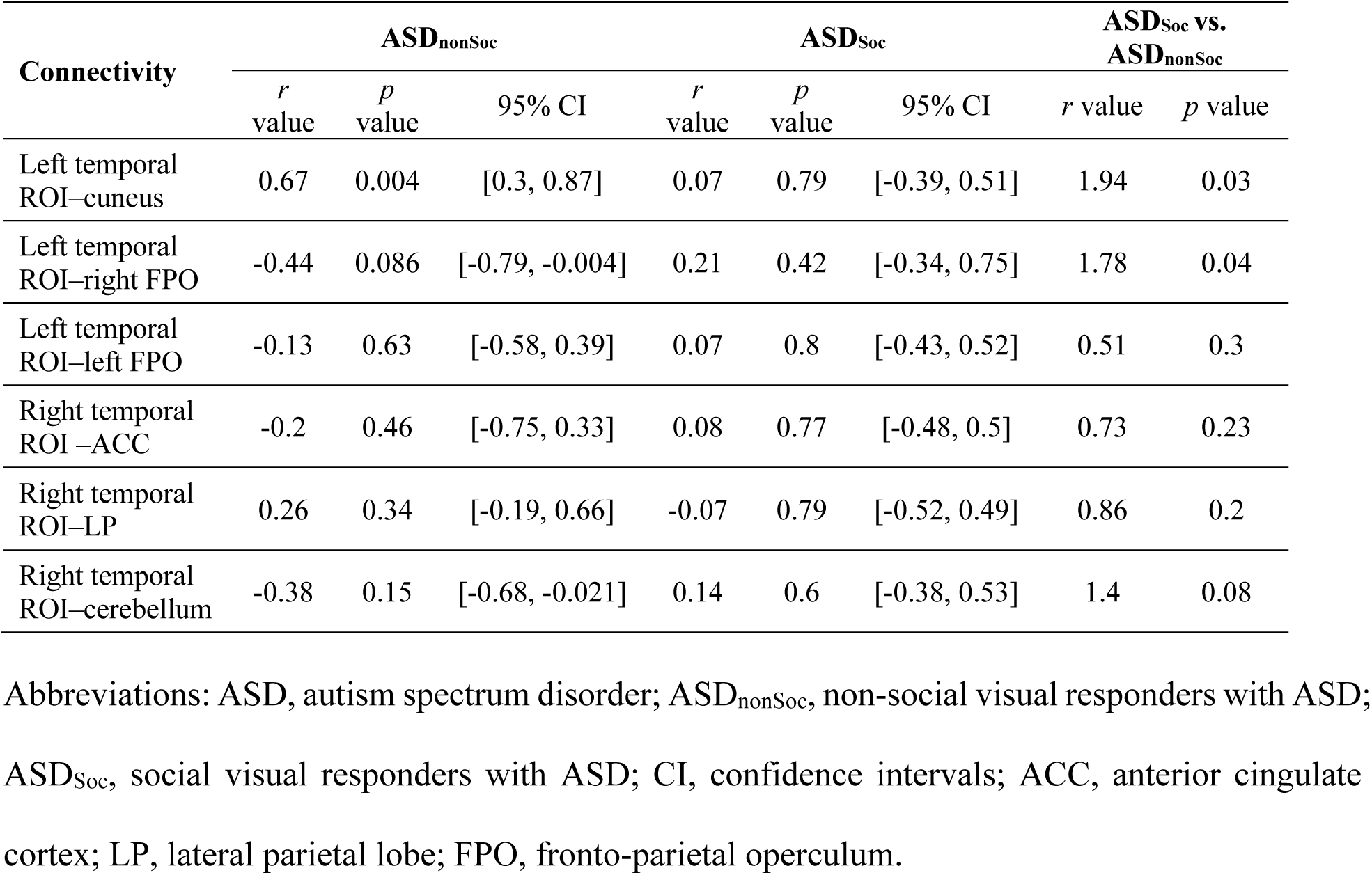
Subgroup–specific associations between functional connectivity and ASD toddler’s social visual attention as well as the comparisons between ASD subgroups.

| Connectivity | ASD<sub>nonSoc</sub> |  |  | ASD<sub>Soc</sub> |  |  | ASD<sub>Soc</sub> vs. ASD<sub>nonSoc</sub> |  |
| --- | --- | --- | --- | --- | --- | --- | --- | --- |
|  | *r* value | *p* value | 95% CI | *r* value | *p* value | 95% CI | *r* value | *p* value |
| Left temporal ROI–cuneus | 0.67 | 0.004 | [0.3, 0.87] | 0.07 | 0.79 | [-0.39, 0.51] | 1.94 | 0.03 |
| Left temporal ROI–right FPO | -0.44 | 0.086 | [-0.79, -0.004] | 0.21 | 0.42 | [-0.34, 0.75] | 1.78 | 0.04 |
| Left temporal ROI–left FPO | -0.13 | 0.63 | [-0.58, 0.39] | 0.07 | 0.8 | [-0.43, 0.52] | 0.51 | 0.3 |
| Right temporal ROI –ACC | -0.2 | 0.46 | [-0.75, 0.33] | 0.08 | 0.77 | [-0.48, 0.5] | 0.73 | 0.23 |
| Right temporal ROI–LP | 0.26 | 0.34 | [-0.19, 0.66] | -0.07 | 0.79 | [-0.52, 0.49] | 0.86 | 0.2 |
| Right temporal ROI–cerebellum | -0.38 | 0.15 | [-0.68, -0.021] | 0.14 | 0.6 | [-0.38, 0.53] | 1.4 | 0.08 |

Abbreviations: ASD, autism spectrum disorder; ASD<sub>nonSoc</sub>, non-social visual responders with ASD; ASD<sub>Soc</sub>, social visual responders with ASD; CI, confidence intervals; ACC, anterior cingulate cortex; LP, lateral parietal lobe; FPO, fronto-parietal operculum.

**Supplementary Figure 1.**
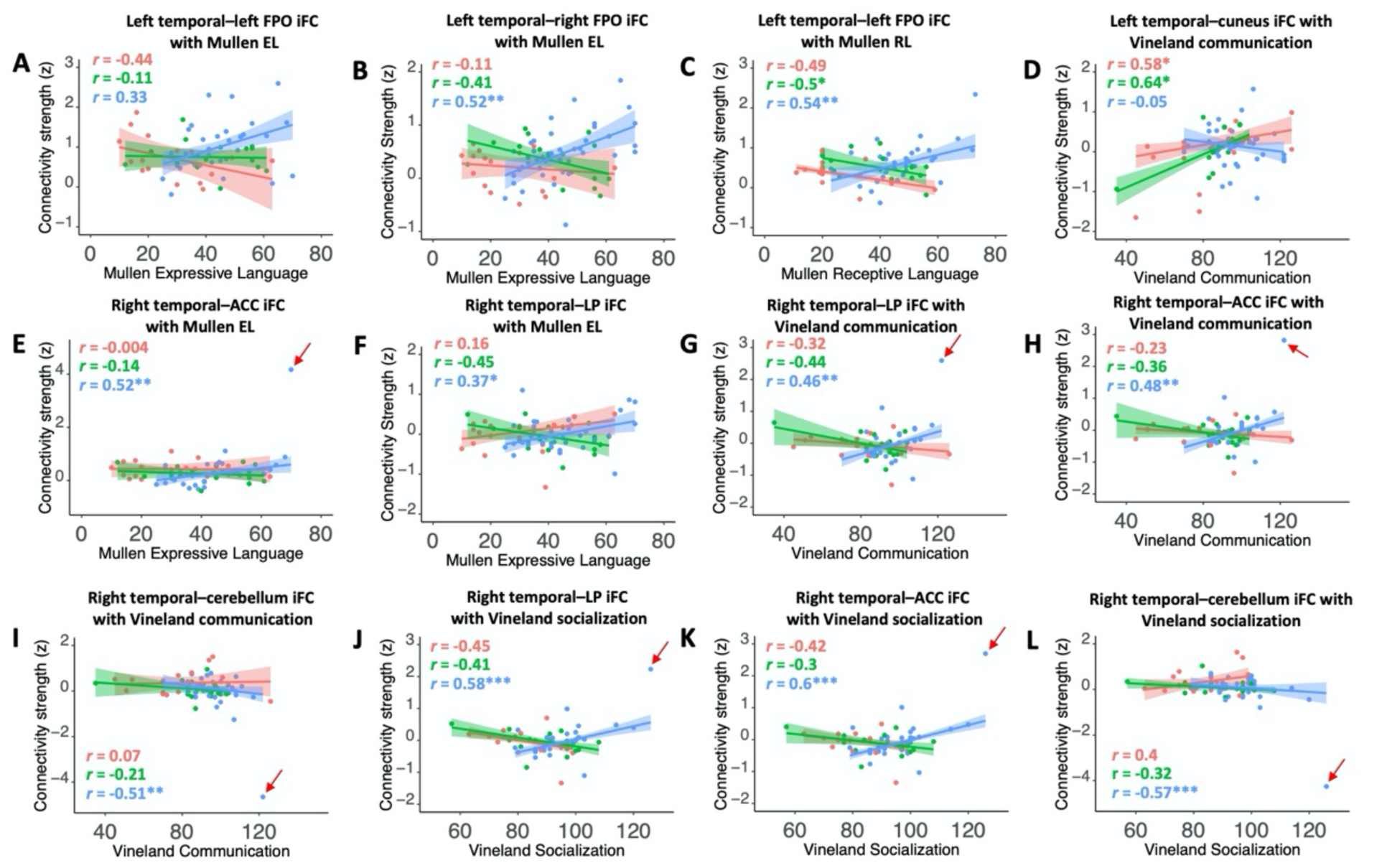
Scatterplots showing connectivity–behavior relationships in ASD subgroups and non-ASD toddlers. Scatterplots display iFC–behavior relationships for: (A) left temporal–left FPO iFC with Mullen expressive language scores; (B) left temporal–right FPO iFC with Mullen expressive language scores; (C) left temporal–left FPO iFC with Mullen receptive language scores; (D) left temporal–cuneus iFC with Vineland socialization scores; (E) right temporal–ACC iFC with Mullen expressive language scores (excluding the outlier in non-ASD: *r*(29) = 0.43; *p* = 0.015); (F) right temporal–LP iFC with Mullen expressive language scores; (G) right temporal–LP iFC with Vineland communication scores (excluding the outlier in non-ASD: *r*(29) = 0.23; *p* = 0.22); (H) right temporal–ACC iFC with Vineland communication scores (excluding the outlier in non-ASD: *r*(29) = 0.25; *p* = 0.17); (I) right temporal–cerebellum iFC with Vineland communication scores (excluding the outlier in non-ASD: *r*(29) = −0.3; *p* = 0.096); (J) right temporal–LP iFC with Vineland socialization scores (excluding the outlier in non-ASD: *r*(29) = 0.33; *p* = 0.069); (K) right temporal–ACC iFC with Vineland socialization scores (excluding the outlier in non-ASD: *r*(29) = 0.327; *p* = 0.073); (L) right temporal–cerebellum iFC with Vineland socialization scores *r(*29) = −0.22; *p* = 0.23). The relationships for ASD_nonSoc_ is shown in pink, for ASD_Soc_ shown in green, and for non-ASD shown in blue. The red arrows point to outliers. * *p* < .05, ** *p* < .005, *** *p* < .001. Abbreviations: iFC, intrinsic functional connectivity; EL, expressive language; RL, receptive language; FPO, fronto-parietal operculum; ACC, anterior cingulate cortex; LP, lateral parietal lobe.

